# Alternative oxidase alleviates mitochondrial oxidative stress during limited nitrate reduction in *Arabidopsis thaliana*

**DOI:** 10.1101/2024.04.20.590391

**Authors:** Daisuke Otomaru, Natsumi Ooi, Kota Monden, Takamasa Suzuki, Ko Noguchi, Tsuyoshi Nakagawa, Takushi Hachiya

## Abstract

The conversion of nitrate to ammonium, i.e., nitrate reduction, is a major consumer of reductants in plants. Previous studies have reported that the mitochondrial alternative oxidase (AOX) is upregulated under limited nitrate reduction conditions, including no/low nitrate or when ammonium is the sole nitrogen (N) source. Electron transfer from ubiquinone to AOX bypasses the proton-pumping complexes III and IV, thereby consuming reductants efficiently. Thus, upregulated AOX under limited nitrate reduction may dissipate excessive reductants and thereby attenuate oxidative stress. Nevertheless, so far there is no firm evidence for this hypothesis due to the lack of experimental systems to analyze the direct relationship between nitrate reduction and AOX. We therefore developed a novel culturing system for *A. thaliana* that manipulates activities of nitrate reduction and AOX separately without causing N starvation, ammonium toxicity, and lack of nitrate signal. Using this system we examined genome-wide gene expression and plant growth to better understand the relationship between AOX and nitrate reduction. Here we show that AOX alleviates mitochondrial oxidative stress and sustains plant growth under limited nitrate reduction.

## 1 | INTRODUCTION

Plants use nitrate and ammonium as main sources of nitrogen (N). The conversion of nitrate to ammonium, i.e., nitrate reduction, requires eight moles of electrons per mole of nitrate, which accounts for about half of the energy required for protein synthesis from nitrate (Hachiya et al., 2007). Thus, ammonium is an energetically superior N source, but most crops prefer nitrate (Miller & Cramer, 2004). In herbaceous plants, nitrate reduction generally occurs in the shoots (Andrews et al., 1992; Hachiya et al., 2020; Scheurwater et al., 2002), and acts as a major sink of reductants generated by the chloroplasts (Rasmusson et al., 2020). Hence, when the nitrate supply is limited, excessive reductants may accumulate within illuminated photosynthetic cells. Recent studies have shown that the NAD(P)H/NAD(P)^+^ ratio and the levels of hydrogen peroxide are increased in *A. thaliana* leaves of ammonium-grown plants compared with nitrate-grown plants (Podgórska et al., 2013; Podgórska et al., 2020; Rasmusson et al, 2020). Moreover, N starvation treatments have been found to enlarge the ascorbate pool in several plant species (Iwagami et al., 2022). Taken together, these findings suggest that a decrease in nitrate reduction could lead to oxidative stress in plants.

It is widely accepted that nitrate reduction significantly alters the action of the plant mitochondrial electron transport chain (mETC) (Rasmusson et al., 2020). In fact, the amount of electrons consumed in nitrate reduction appears to be comparable to that in the mETC (Noctor & Foyer, 1998; Rasmusson et al., 2020). In photosynthetic tissues, the expression and activities of several enzymes that bypass energy conservation steps in mETC are induced under limited nitrate reduction conditions; i.e., if the plant is experiencing N starvation or if ammonium is the sole N source (Escobar et al., 2006; Hachiya et al., 2010; Noguchi & Terashima, 2006; Qiao et al., 2022; Watanabe et al., 2010). One enzyme is the alternative oxidase (AOX), which allows direct electron transfer from ubiquinone to molecular oxygen, bypassing the proton-pumping complexes III and IV (Noguchi & Yoshida, 2008). Therefore, upregulation of AOX under limited nitrate reduction may dissipate excessive reductants without being limited to steep proton gradients across the mitochondrial inner membrane, thereby attenuating reactive oxygen species (ROS) production and oxidative stress. Actually, in *Arabidopsis* plants under low nitrate conditions, the shoot expression of antioxidant enzyme genes was induced by disrupting the major isoform *AOX1a* (Watanabe et al., 2010). Antisense suppression of *Arabidopsis AOX1a* was found to greatly decrease the reducing state of ascorbate under ammonium but not under nitrate (Podgórska et al., 2020). In barley grown under low nitrate, chemical inhibition of AOX activity elevated the levels of hydrogen peroxide (Wang et al., 2016). Moreover, in tobacco cell cultures subjected to N starvation, antisense suppression of *AOX1* caused carbohydrate accumulation (Sieger et al., 2005). This implies that, under limited nitrate reduction, oxidation of NADH via AOX could replenish the NAD^+^ to drive the glycolysis.

The above studies suggest that AOX is linked to nitrate reduction. However, these studies manipulated nitrate reduction activity by transferring plants to the growth conditions that included no/low nitrate or the use of ammonium as the sole N source. Reduced nitrate reduction may therefore be accompanied by N starvation or ammonium toxicity (Hachiya et al., 2021a), causing plant growth suppression and/or the initiation of stress responses. Moreover, since nitrate acts as a signal to alter genome-wide gene expression (Liu et al., 2017; Wang et al., 2004), a decrease in the nitrate supply also depletes the nitrate signal. For these reasons, it is impossible to distinguish whether the above-mentioned effects of AOX deficiency are caused by reduced nitrate reduction, N starvation, ammonium toxicity, or the lack of nitrate signal. To solve this problem, we developed a novel culturing system to manipulate the degree of nitrate reduction and AOX activity without causing N starvation, ammonium toxicity, and the lack of nitrate signal. Using this system we examined genome-wide gene expression and plant growth to better understand the relationship between AOX and nitrate reduction.

## 2 | MATERIALS AND METHODS

### 2.1 Plant materials and growth conditions

In this study we used *A. thaliana*, accession Columbia (Col-0) as a control line as well as the *aox1a-1* (SALK_084897, Giraud et al., 2008), *aox1a-2* (SAIL_030_D08, Giraud et al., 2008), *nr* (*nia1-1/chl3-5*, Wang et al., 2004), *aox1a-1 nr*, and *aox1a-2 nr* mutants. The homozygous multiple mutants of *aox1a-1 nr* and *aox1a-2 nr* were produced via crossing. The surface-sterilized seeds were placed in the dark at 4°C for three days to break seed dormancy. For *in vitro* culturing, we first grew plants on a sterile cellophane sheet (Hachiya et al., 2021b) placed on a solid medium containing 30 mL half-strength Murashige and Skoog (1/2-MS) salts without nitrogen supplemented with 2.5 mM (NH_4_)_2_SO_4_, 0.1% (w/v) MES-KOH (pH 6.7), 0.5% (w/v) sucrose, and 0.25% (w/v) gellan gum (Fujifilm Wako, Osaka, Japan). Plants were grown horizontally at 22°C for 18 days under a fixed photosynthetic photon flux density (PPFD) of 30 µmol m^−2^ s^−1^ (i.e., constant light). Plants-on-cellophane were then transferred to a medium containing 30 mL of 1/2-MS salts without nitrogen supplemented with 0.05% (w/v) MES-KOH (pH 5.7), 0.5% (w/v) sucrose, and 0.25% (w/v) gellan gum (Fujifilm). The resulting mixture was incubated at 22°C for 24 h, also under a PPFD of 30 µmol m^−2^ s^−1^ (i.e., constant light). Finally, the plants-on-cellophane were transferred to a medium containing 30 mL of 1/2-MS salts supplemented with 0.05% (w/v) MES-KOH (pH 5.7), 0.5% (w/v) sucrose, and 0.25% (w/v) gellan gum (Fujifilm) and then incubated at 22°C under a PPFD of 150–200 µmol m^−2^ s^−1^ (constant light). For soil cultivation, plants were grown for 14 or 24 days under a PPFD of 100–130 µmol m^−2^ s^−1^ (i.e., on a 16h/8h light/dark cycle) at 22–23°C using an Arasystem 360 kit (Betatech BVBA, Gent, Belgium) in a 1: 1 mixture of nutrient-rich soil (Supermix A; Sakata Seed, Kyoto, Japan) and vermiculite.

### 2.2 Determination of in vitro nitrate reductase activities

First, we assessed *in vitro* nitrate reductase (NR) activity using a previously published method (Hachiya et al., 2016) with slight modification. Frozen samples were first ground in a Multi-Bead Shocker (Yasui Kikai, Osaka, Japan) using zirconia beads. The resulting powder was then mixed with 10 vol. of extraction buffer (50 mM HEPES-KOH, pH 7.6, 1 mM EDTA, 7 mM cysteine) and the extracts were centrifuged at 20,400×g at 4°C for 10 min. Next, a 30 μl aliquot of the supernatant was added to 90 μl of assay buffer (50 mM HEPES-KOH, pH 7.6, 133 μM NADH, 2 mM EDTA, 6.67 mM KNO_3_). After incubation at 30°C for 2 and 17 min, 24 μl of 1% (w/v) sulfanilamide solution in 1 N HCl and 24 µl of 0.02% (w/v) *N*-(1-naphthyl)ethylenediamine dihydrochloride solution were added to 48 µl aliquots of reaction mixture. Finally, nitrite content was determined based on absorbance readings at 540 nm. *In vitro* NR activity was calculated by determining the nitrite amount produced over a 15 min period.

### 2.3 Determination of in vivo nitrate reductase activities

First, we submerged fresh shoots in 50 vols. of reaction buffer (i.e., 100 mM sodium phosphate buffer, pH 7.4, 10 mM KNO_3_, 4% (v/v) n-propanol), followed by vacuum-infiltration. The reaction mixture was then incubated at 30°C for 1 h in the dark. The amount of nitrate produced was then visualized by mixing the supernatant with 1% (w/v) sulfanilamide solution in 1 N HCl, and 0.02% (w/v) *N*-(1-naphthyl) ethylenediamine dihydrochloride solution in a 2: 1: 1 ratio.

### 2.4 Determination of nitrate concentration

Nitrate content was quantified as per a previously published protocol (Hachiya et al., 2016) with slight modifications. Nitrate was first extracted with 10 vol. of deionized water at 100°C for 20 min. Next, 10 µL of the supernatant was mixed with 40 µL of a reaction reagent (i.e., 5% (w/v) salicylic acid in concentrated sulfuric acid), and the resulting mixture was then incubated at room temperature for 20 min. A mock treatment of 40 µL of concentrated sulfuric acid was also produced. Finally, 1 mL of 8% (w/v) NaOH solution was added to the mixture, and nitrate was determined based on absorbance at 410 nm.

### 2.5 Determination of total protein

Total protein was determined according to a previously described method (Hachiya et al., 2016). Frozen samples were first homogenized with a Multi-Beads Shocker (Yasui Kikai) using zirconia beads. Total proteins were then extracted with 10 vol. of sample buffer [2% (w/v) SDS, 62.5 mM Tris-HCl (pH 6.8), 10% (v/v) glycerol, and 0.0125% (w/v) bromophenol blue] and Halt^TM^ protease inhibitor cocktail (ThermoFisher Scientific, Tokyo, Japan), followed by incubation at 95°C for 5 min. Extracts were then centrifuged at 20,400×g at 8°C for 10 min and 10 μL aliquots were suspended in 500 μL deionized water. Next, 100 μL 0.15% (w/v) sodium deoxycholate was added, and the mixture was incubated at room temperature for 10 min. 100 μL of 72% (v/v) trichloroacetic acid was then added, followed by incubation at room temperature for 15 min and centrifugation at 20,400×g for 10 min. Precipitates were then air-dried and suspended in 25 µL water. This suspension was used to determine total protein concentrations using Takara BCA Protein Assay Kits (TaKaRa, Kusatsu, Japan).

### 2.6 Immunodetection of AOX and ACTIN proteins

12.35 µL of total protein extract was mixed with 0.65 µl of 1 M DTT, followed by incubation at 95°C for 3 min. 5 µL of mixture (∼0.43 mg fresh weight) was then subjected to SDS-PAGE using a 12% Mini-PROTEAN TGX Gel (Bio-Rad, Hercules, CA, USA) before transferring to a Trans-Blot Turbo Mini PVDF membrane using the Trans-Blot Turbo Transfer System (Bio-Rad). Membranes were incubated for 1h in blocking buffer (50 mM Tris-HCl, and 150 mM NaCl, pH 7.6, 0.1% (v/v) Tween-20) containing 5% (w/v) ECL Prime Blocking Agent (Cytiva, Tokyo, Japan). Membranes were then incubated for 1h with a 1/200 dilution of monoclonal antibodies against AOX (AS10 699, Agrisera, Vännäs, Sweden) or a 1/2,500 dilution of polyclonal antibodies against ACTIN (AS13 2640, Agrisera). Finally, the antigen–antibody complex was detected using a 1/10,000 dilution of horseradish peroxidase conjugated with anti-mouse IgG sheep antibodies (Cytiva) for AOX or anti-rabbit IgG donkey antibodies (Cytiva) for ACTIN by ECL Prime chemiluminescent detection (Cytiva). Proteins blotted on membranes were visualized using TaKaRa CBB Protein Safe Stain (TaKaRa) and chemiluminescence and CBB staining were visualized using ImageQuant LAS 500 mini (Cytiva).

### 2.7 RNA extraction

Samples were first homogenized as described above. Next, total RNA was extracted using an RNeasy Plant Mini Kit (Qiagen, Tokyo, Japan). For RNA-seq library preparation, RNA was purified using on-column DNase digestion (Qiagen).

### 2.8 RNA-seq

RNA quality was first evaluated using a Qubit RNA IQ Assay Kit (ThermoFisher Scientific). RNA samples with RNA IQs 8.7–10.0 were used for library preparation. cDNA libraries were constructed using a NEBNext Ultra II RNA Library Prep Kit with Sample Purification Beads (New England Biolabs, Tokyo, Japan), a NEBNext Poly(A) mRNA Magnetic Isolation Module (New England Biolabs), and NEBNext Multiplex Oligos for Illumina (New England Biolabs). cDNA libraries were then sequenced using a NextSeq 500 (Illumina, Tokyo, Japan), and resulting bcl files were converted to fastq files using bcl2fastq (Illumina). The RNA-seq raw data are available in ArrayExpress database under accession number E-MTAB-14027. Finally, obtained reads were analyzed using iDEP version 096 (Ge et al., 2018), Metascape (Zhou et al., 2019), and GeneCloud (Krouk et al., 2015) using the default settings.

### 2.9 RT-qPCR

Reverse transcription (RT) was performed using a ReverTraAce qPCR RT Master Mix with gDNA Remover (Toyobo, Osaka Japan). Synthesized cDNA was then diluted tenfold with water and used for quantitative PCR (qPCR). RT-qPCR was performed on a QuantStudio 1 (ThermoFisher Scientific) with KOD SYBR qPCR Mix (Toyobo). Relative transcript levels were calculated using the comparative cycle threshold method with *ACTIN3* as an internal standard. Primer sequences are shown in Table S1.

### 2.10. Statistical analysis

The Tukey–Kramer multiple comparison test were conducted using R software v.2.15.3.

## 3 | RESULTS and DISCUSSION

### 3.1 Manipulation of nitrate reduction and AOX activities

We manipulated the activities of nitrate reduction and AOX via the transfer experiment (Figure 1A) using Col-0 and the mutants deficient in either or both of NR and AOX activities. First, plants were grown for 18 days with ammonium as the sole N source to ensure uniform growth independently of NR activity. Higher medium pH and lower light intensity were used to reduce ammonium toxicity and AOX expression (Escobar et al., 2006; Hachiya et al., 2010; Hachiya et al., 2021a; Yoshida et al., 2011), allowing uniform growth regardless of AOX activity. Second, plants were subjected to N starvation at pH 5.7 for 24 h to induce AOX expression (Escobar et al., 2006; Hachiya et al., 2010). Finally, plants were transferred to a medium containing adequate nitrate and incubated at an elevated light intensity, which induces NR expression and fills photosynthetically-derived reductants to the cell (Noguchi & Yoshida, 2008; Wang et al., 2004; Yoshida et al., 2010). The NR activity of Col-0 increased rapidly from 0 to 7 h after nitrate supply and slightly more from 7 to 24 h (Figure S1A). Both *in vitro* and *in vivo* NR activities were increased 7 h after nitrate supply in the shoots of Col-0 and *aox1a-1*, but not in *nr* and *aox1a-1 nr* (Figures 1B and 1C). Thus, we focused on the 7 h after nitrate supply to identify the early effects of nitrate reduction for subsequent experiments. We found that nitrate accumulated in the shoots of all lines after nitrate supply, with concentrations ranging from 14.7 to 21.6 µmol g^−1^ (Figure 1D), indicating an adequate supply of nitrate for signaling. Actually, expression of *NIA2* encoding the major NR isoform in leaves, which is inducible by nitrate signal (Konishi & Yanagisawa, 2013; Wilkinson & Crawford, 1991), was strongly induced 7 h after nitrate supply (Figure S1B). Higher nitrate concentrations in shoots of *nr* and *aox1a-1 nr* than Col-0 and *aox1a-1* would reflect a deficiency in nitrate reduction (Figure 1D). Shoot protein concentrations were comparable among all lines before and after nitrate supply, ranging from 27.0 to 29.7 mg g^−1^ (Figure 1E), suggesting that no N starvation occurred. RT-qPCR and Western blot analyses confirmed that the signals corresponding to *AOX1a* and AOX were negligible in *aox1a-1* and *aox1a-1 nr* (Figures S1C and S1D), suggesting that knockout of *AOX1a* is sufficient to diminish AOX activity, as reported earlier (Giraud et al., 2008; Hachiya et al., 2010). Taken together, these culturing conditions permit comparisons of plants differing nitrate reduction and AOX activity without causing ammonium toxicity, N starvation, and lack of nitrate signaling.

**FIGURE 1.**
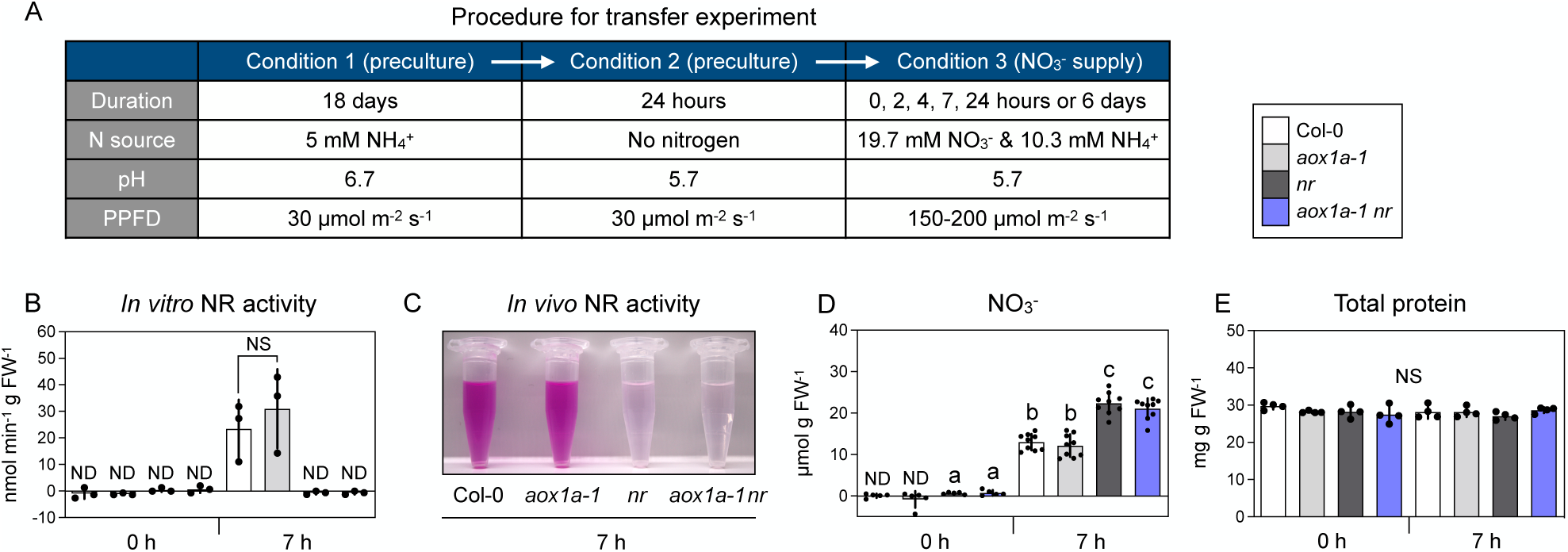
Manipulation of activities of nitrate reduction and AOX without causing N starvation, ammonium toxicity, and the lack of nitrate signal. (A) Schematic diagram of experiment. (B-E) *in vitro* NR activity (B), *in vivo* NR activity (C), nitrate concentration (D), and total protein concentration (E) in plant shoots before and 7 h after nitrate supply. Two plants of each line (eight in total) per plate were grown, and two shoots were pooled as one biological replicate. Data: mean ± SD (n = 3 (B), n = 5–9 (D), n = 4 (E)). For (C), color intensity is proportional to NR activity. Different lowercase letters indicate significant differences determined via Tukey–Kramer tests at *P* < 0.05. ND and NS denote “not detected” and “not significant.” FW denotes “fresh weight.”

### 3.2 AOX alleviates mitochondrial oxidative stress under limited nitrate reduction

Next, to dissect the role of AOX in limited nitrate reduction, we performed three independent RNA-seq analyses using shoots from Col-0, *aox1a-1*, *aox1a-2*, *nr*, *aox1a-1 nr*, and *aox1a-2 nr* 7 h after nitrate supply (Table S2). A k-means clustering analysis classified transcripts into four groups according to their expression patterns (Figure 2A and Table S3). In cluster D, *AOX1a* deficiency consistently induced gene expression in the *nr* background, but not in the Col-0 background (Figures 2A and 2B). Moreover, 42 of 141 genes in cluster D were statistically significantly upregulated in *aox1a-1 nr* and *aox1a-2 nr* relative to *nr* (Figure 2C and Table S4). Further enrichment analyses of the 42 genes identified significant overrepresentation of the terms “toxin catabolic process,” “glutathione metabolism” (Figure 2D), “interpro-ipr004046 (glutathione S-transferase, C-terminal),” and “gst” (Figure 2E). These terms were derived from the glutathione S-transferase (GST) genes *AT2G29460* (*GSTU4*), *AT1G17170* (*GSTU24*), *AT1G17180* (*GSTU25*), and *AT1G02920* (*GSTF7*) (Table S4). *GSTU4*, a plant-specific tau class GST, may contribute to hydrogen peroxide degradation by using glutathione as an electron donor (Santamaría et al., 2018). The terms “response to hypoxia” and “response to oxidative stress” were also enriched in these 42 genes (Figure 2D). Furthermore, genes induced by hypoxia (Tamura & Bono, 2022) and H_2_O_2_ treatments (Hieno et al., 2019) were significantly upregulated in *nr*, and their induction was enhanced by *AOX1a* deficiency (Figures 2F and 2G and Tables S5 and S6). The NAC transcription factor ANAC017 mediates ROS-related mitochondrial retrograde signaling, thereby activating mitochondrial dysfunction stimulation genes including *AOX1a*, *UPOX*, and *ANAC013* (De Clercq et al., 2013; Ng et al., 2013). The expression of ANAC017-inducible genes (Ng et al., 2013) and mitochondrial dysfunction stimulation genes (De Clercq et al., 2013) reached a maximum level in *aox1a nr and aox1a-2 nr* (Figures 2H and 2I and Tables S7 and S8). Further RT-qPCR analyses revealed that the expression levels of hypoxia-inducible genes (Figures 2J–M) and oxidative stress marker genes (Figures 2N–Q) were induced by nitrate supply in *nr*, and this induction was enhanced by *AOX1a* deficiency. Meanwhile, before nitrate supply, little difference was observed among all lines (Figures 2J–Q). Since hypoxia elevates intracellular NADH/NAD^+^ ratio via mETC inhibition (Igamberdiev et al., 2005), the upregulation of hypoxia-inducible genes may reflect reductant accumulation. Consequently, our transcriptome analysis suggests that AOX dissipates excessive reductants and mitigate oxidative stress under limited nitrate reduction.

**FIGURE 2.**
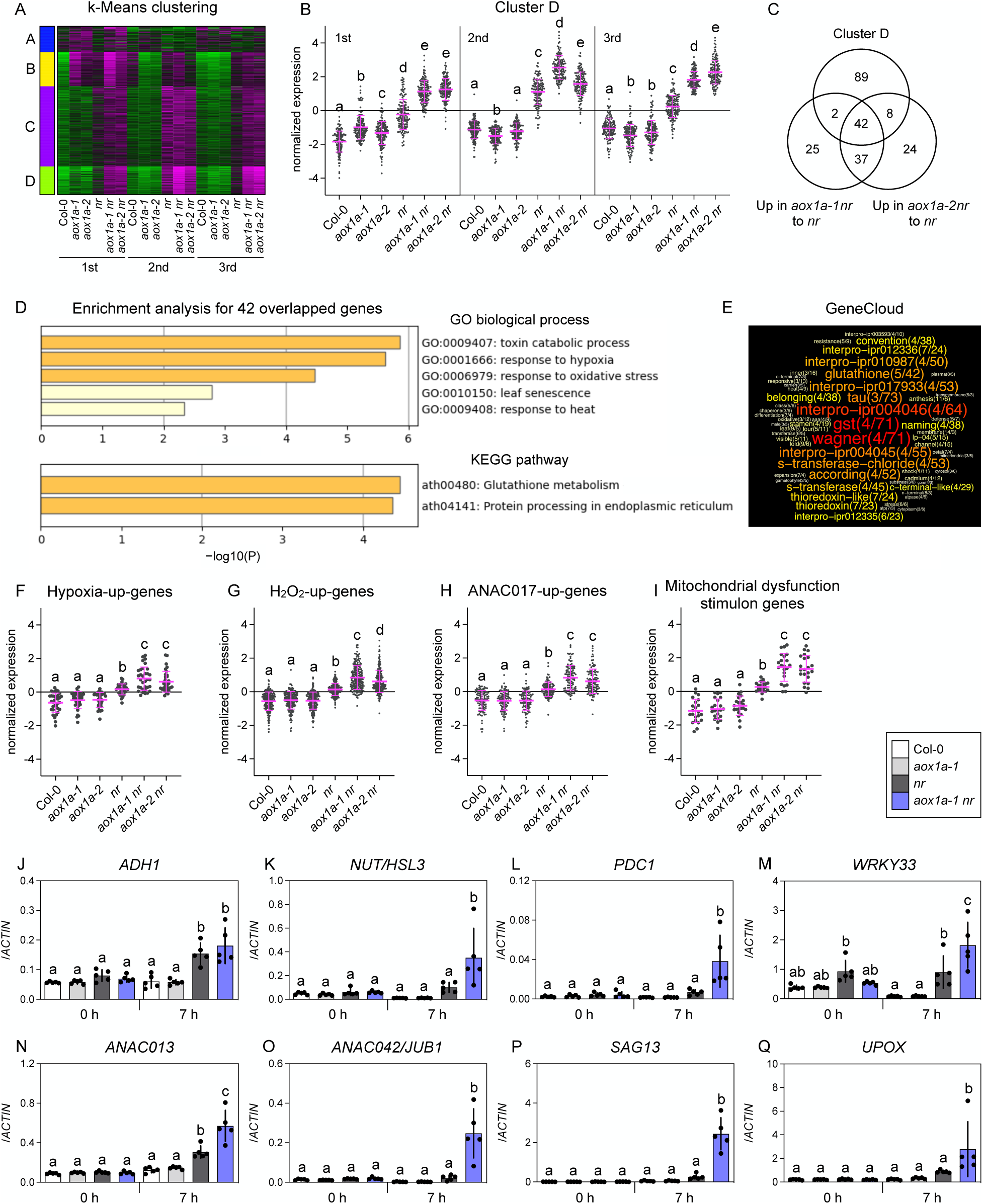
Transcriptomic alteration by *AOX1a* deficiency under limited nitrate reduction conditions. (A–I) One plant from each line (six in total) per plate was grown. Six shoots were pooled as one biological replicate. Shoots from plants 7 h after nitrate supply were subjected to RNA-seq. Numbers indicate the order of RNA-seq experiments. (A) Heat map from a k-means clustering of normalized transcript levels. Magenta and green represent higher and lower expression levels, respectively. (B) Plots of normalized transcript levels in cluster D. Normalized transcript levels of splice variants were averaged for each gene. (C) Venn diagram showing gene number from cluster D and genes significantly upregulated in *aox1a-1 nr* or *aox1a-2 nr* relative to *nr*. (D, E) Enriched terms identified by Metascape (D) and GeneCloud (E) from the 42 genes in (C). (E) shows the number of genes containing the term (left) and the fold enrichment (right). (F–I) Shoot expression of genes upregulated under hypoxia (F), by H_2_O_2_ (G), by ANAC017 (H), and included in mitochondrial dysfunction stimulon (I) 7h after nitrate supply. Normalized transcript levels of splice variants were averaged for each gene using means from three independent experiments. (J– Q) RT-qPCR analysis of marker genes upregulated under hypoxia (J–M) and by oxidative stress (N–Q) in shoots before and 7 h after nitrate supply. Two plants of each line (eight in total) per plate were grown, and four shoots were pooled as one biological replicate. Data: mean ± SD (n = 5). (B, F–Q) Different lowercase letters indicate significant differences determined via Tukey– Kramer tests at *P* < 0.05.

The plant mETC possesses type II NAD(P)H dehydrogenases located on the cytosol side (ND_ex_) or matrix side (ND_in_) of the inner mitochondrial membrane (Rasmusson et al., 2020). Since these dehydrogenases transfer electrons to ubiquinone and bypass the proton-pumping complex I, they can dissipate excessive reductants like AOX. Of these, the expression of *NDB2*, encoding the primary ND_ex_ contributor to NADH oxidation, was significantly induced in *nr* after nitrate supply, and this induction was intensified by *AOX1a* deficiency (Figures 3A and 3B). Since cytosolic NR has a much lower K_m_ for NADH than ND_ex_ (Sweetman et al., 2019), NDB2 may operate only when nitrate reduction is limited. The *NDA2*, encoding the NADH-oxidizing ND_in_ (Rasmusson et al., 2020), also showed a expression pattern similar to *NDB2* after nitrate supply (Fig. 3A and 3C). The compensated induction of *NDB2* and *NDA2* by *AOX1a* deficiency (Figures 3) suggests a contribution of AOX1a-NDB2 and AOX1a-NDA2 modules to balance cellular redox under limited nitrate reduction. Also, the STRING database ver. 12.0 integrating protein-protein interactions (Szklarczyk et al., 2023) confirmed tight functional connections between AOX1a, NDB2, and NDA2. Meanwhile, the expression of the uncoupling proteins (UCPs), which act as an uncoupler and/or as an aspartate/glutamate exchanger across the inner mitochondrial membrane (Monné et al., 2018), was little changed among the lines (Figure 3A).

**FIGURE 3.**
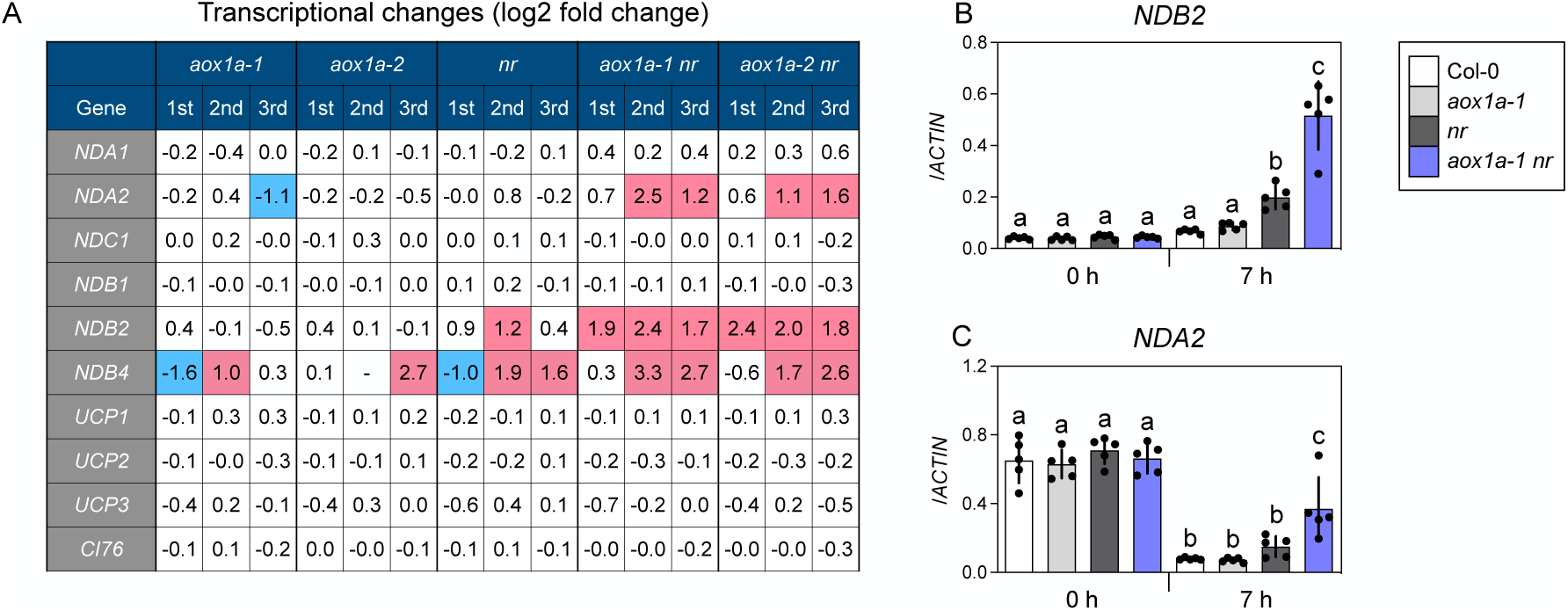
Effects of *AOX1a* deficiency on gene expression of type II NADP(H) dehydrogenases and uncoupling proteins under limited nitrate reduction conditions. (A) Transcriptional changes in genes encoding type II NADP(H) dehydrogenases (*NDA1/2*, *NDB1/2/4*, *NDC1*), uncoupling proteins (*UCP1/2/3*), and *CI76* encoding the complex I 76-kD subunit in plant shoots 7h after nitrate supply. Transcriptional changes are represented as log2 fold change ratios against Col-0 based on the RPM (reads per million mapped reads) from the RNA-seq. Red and blue represent up-(>2-fold) and downregulation (<0.5 fold), respectively. *NDB3* was removed due to low expression. (B,C) Relative transcript levels of *NDB2* (B) and *NDA2* (C) in plant shoots before and 7 h after nitrate supply. Data: mean ± SD (n = 5). Different lowercase letters indicate significant differences determined via Tukey–Kramer tests at *P* < 0.05.

### 3.3 AOX sustains plant growth under limited nitrate reduction

Finally, we analyzed shoot growth parameters following nitrate supply. Over 6 days, we found that *AOX1a* deficiency significantly reduced (−15%) shoot fresh weight in *nr*, but not in Col-0 (Figure 4A). Meanwhile, rosette diameter was little affected by *AOX1a* deficiency (Figure 4B). When plants were grown in nutrient-rich soil, shoot appearance and shoot fresh weight were decreased in *aox1a-1 nr* and *aox1a-2 nr* relative to the others (Figures 4C–E). However, in NR-deficient plants, leaves were pale green (Figure 4C). Thus, it is plausible that long-term limitation of nitrate reduction could cause N starvation, which complicates the interpretation of the relationship between nitrate reduction, AOX, and shoot growth.

**FIGURE 4.**
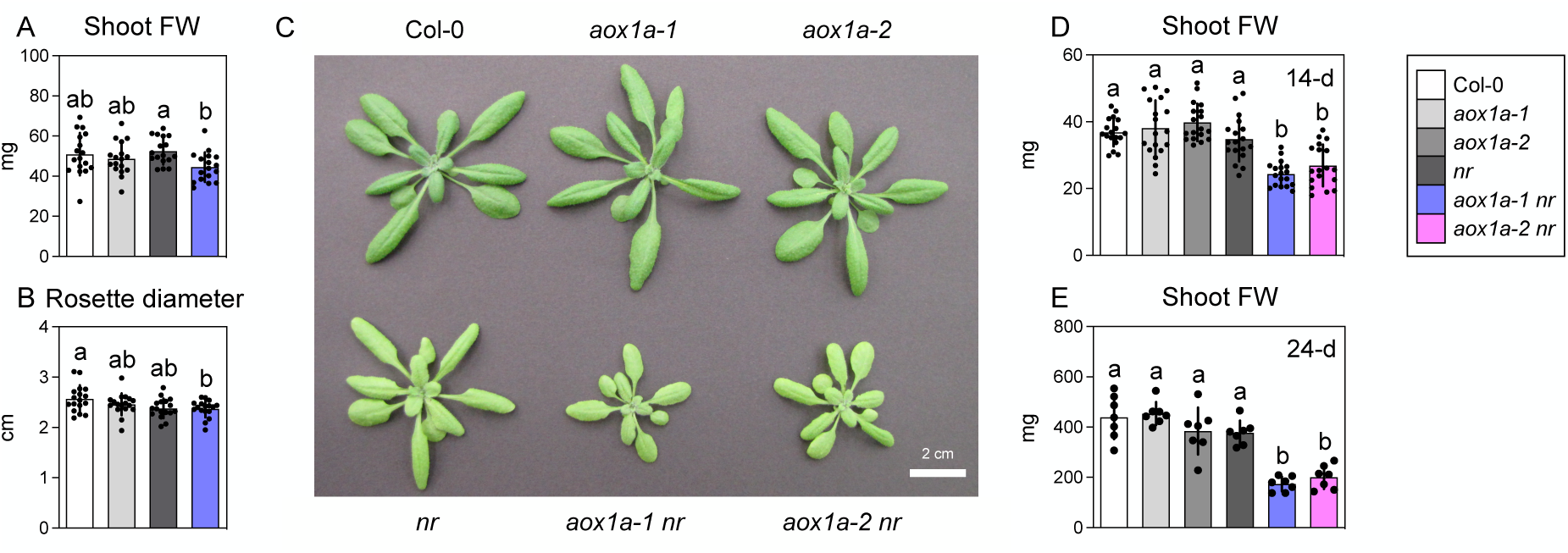
Effects of *AOX1a* deficiency on shoot growth under limited nitrate reduction conditions. (A, B) Shoot fresh weight (A) and rosette diameter (B) of plants 6 days after nitrate supply. One plant of each line (four in total) per plate were grown and one shoot was pooled as one biological replicate. (C–E) Shoot appearance of 24-day-old plants (C) and shoot fresh weights of 14-day-old (D) and 24-day-old (E) plants grown in pots containing nutrient-rich soil. Data: mean ± SD (n = 17 (A, B), n = 18 (D), n = 7 (E)). Different lowercase letters indicate significant differences determined via Tukey–Kramer tests at *P* < 0.05.

## 4 | CONCLUSIONS

We have successfully developed a cultural system to manipulate activities of nitrate reduction and AOX without causing ammonium toxicity, N starvation, and lack of nitrate signaling. Analyses using this system suggest that AOX alleviates mitochondrial oxidative stress and sustain plant growth under limited nitrate reduction. It awaits further analyses to address physiological and biochemical details using the system based on knowledge obtained from the present study.

## AUTHOR CONTRIBUTIONS

DO and NO performed most of the measurements and downstream analyses. KM supported most of the measurements and downstream analyses. TS performed next-generation sequencing analysis. KN and TN supervised all analyses and interpretations. TH designed the study, performed a part of growth analysis, and supervised all analyses and interpretations. All authors contributed to the writing and/or reviewing of the manuscript.

## ACKNOWLEDGEMENTS

*aox1a-2* seeds were provided by Dr. James Whelan (La Trobe University, Australia). The authors thank Enago for English language review.

## DATA AVAILABILITY STATEMENT

The RNA-seq raw data are available in ArrayExpress under accession number E-MTAB-14027 (https://www.ebi.ac.uk/biostudies/arrayexpress).

## SUPPORTING INFORMATION

Additional supporting information can be found online in the Supporting Information section at the end of this article.

## Notes

### Competing Interest Statement

The authors have declared no competing interest.

### Summary of Updates

We have added a few data (Figure 3C, Supplementary Figures 1B and 1C) with the relevant descriptions and updated the format aiming for submitting the manuscript to another journal.

