## SUPPORTING_FIGURE 1 for "Alternative oxidase alleviates mitochondrial oxidative stress during limited nitrate reduction in *Arabidopsis thaliana*"

### Supplementary Figure 1

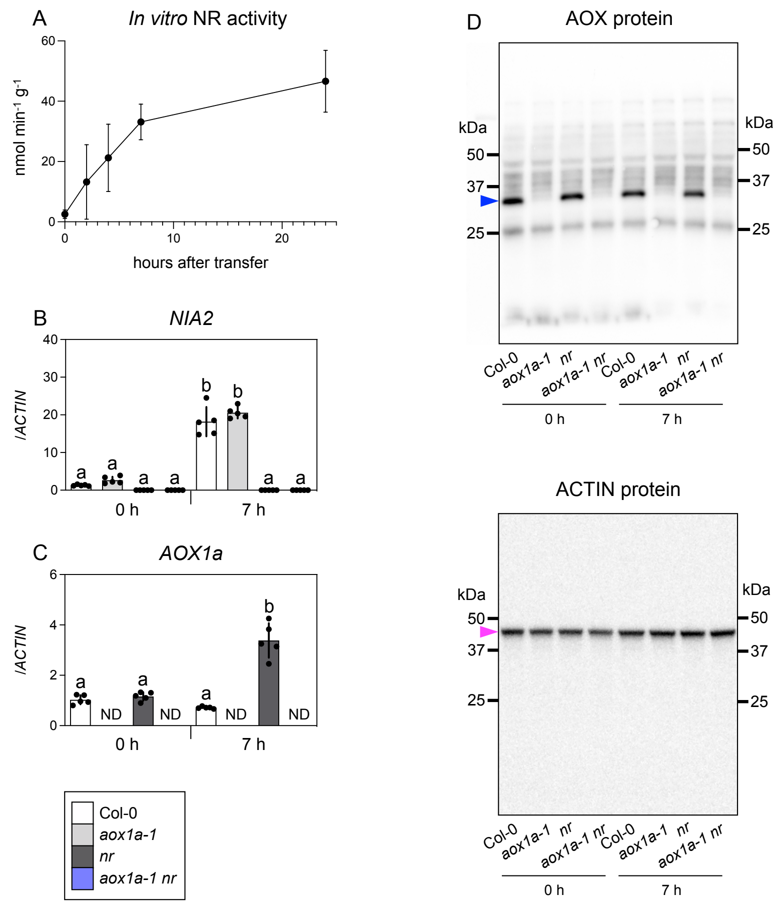


**Fig. S1. Manipulation of activities of nitrate reduction and AOX.**
(A) Time-course of *in vitro* NR activity in Col-0 shoots after nitrate supply. Data: mean ± SD (n = 6). Shoots from eight plants per plate were pooled as one biological replicate. (B,C) RT-qPCR analysis of *NIA2* (B) and *AOX1a* (C) in shoots before and 7 h after nitrate supply. Two plants of each line (eight in total) per plate were grown, and four shoots were pooled as one biological replicate. Data: mean ± SD (n = 5). (D) Immunodetection of AOX and ACTIN isoproteins with specific antisera (Unclopped images). The blue and magenta arrowheads denote the signals corresponding to AOX and ACTIN, respectively.
